## Supplementary Figures for "Distinguishing examples while building concepts in hippocampal and artificial networks"

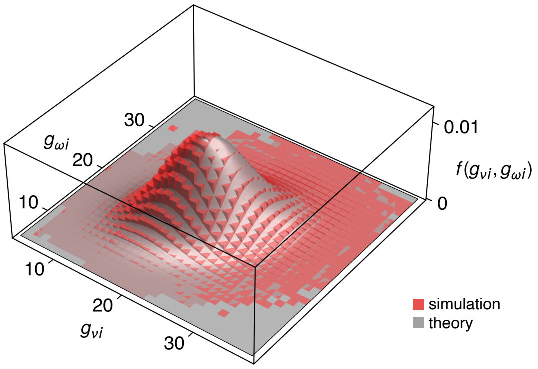


**Figure S1**: Extended results for Supplementary Methods. Joint probability distribution of total inputs $G_{\nu i}$ and $G_{\omega i}$ on postsynaptic neuron $i$ for two patterns $\nu \neq \omega$ (Eq. S12). The theoretically derived probability density function $f(g_{\nu i}, g_{\omega i})$ (Eq. S25) agrees with simulation results. Each $g$ is a sample of the corresponding random variable $G$. Simulation parameters are 1000 presynaptic neurons, 1000 postsynaptic neurons, synaptic connection probability 0.1, 100 random activity patterns, presynaptic pattern density 0.1, and presynaptic correlation 0.3.

*

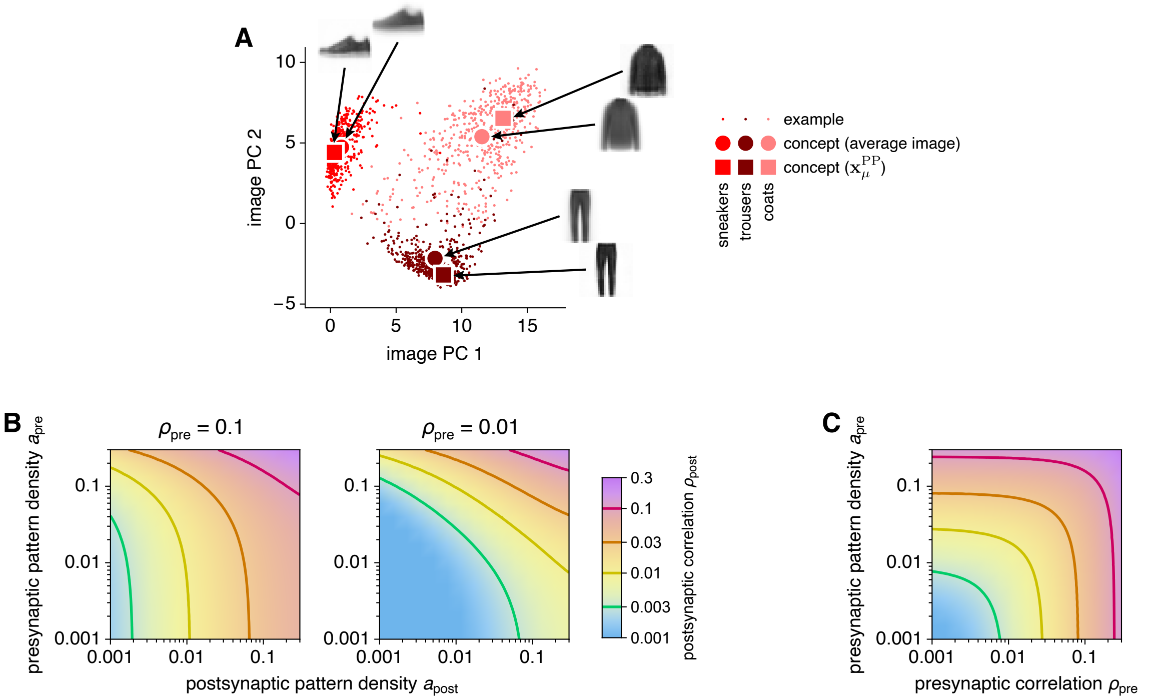


**Figure S2**: Extended results for Fig. 2 of the main text. (**A**) Example images and two definitions of concepts projected along the first two principal components (PCs) of the example dataset. Averaged images are formed by averaging pixel intensities. PP concept target patterns $\mathbf{x}_\mu^{\mathrm{PP}}$ are formed according to Eq. 13 of the main text and then passed through the visualization pathways in Fig. 2F of the main text. (**B**, **C**) Postsynaptic correlation $\rho_{\mathrm{post}}$ in binary feedforward networks in Fig. 2D, E. (**B**) $\rho_{\mathrm{post}}$ as a function of presynaptic pattern density $a_{\mathrm{pre}}$ and postsynaptic pattern density $a_{\mathrm{post}}$ for two values of presynaptic correlation $\rho_{\mathrm{pre}}$. On the left, $\rho_{\mathrm{pre}} = 0.1$ corresponds to the red line, and on the right, $\rho_{\mathrm{pre}} = 0.01$ corresponds to the yellow line. Decorrelation occurs in the regions below and to the left of these lines. (**C**) $\rho_{\mathrm{post}}$ as a function of presynaptic pattern density $a_{\mathrm{pre}}$ and presynaptic correlation $\rho_{\mathrm{pre}}$ for postsynaptic pattern density $a_{\mathrm{post}} = 0.1$.

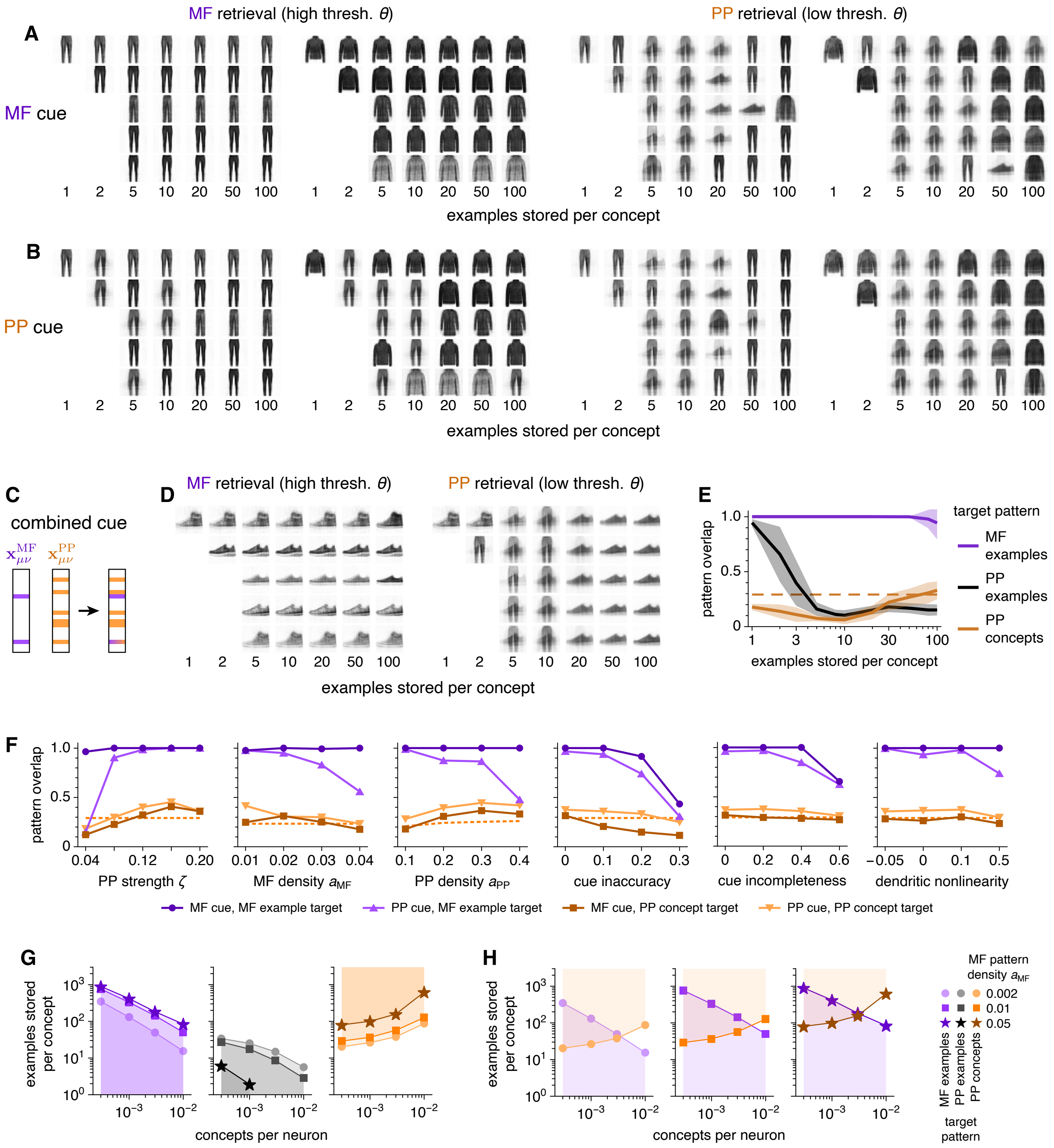


**Figure S3**: Extended results for Fig. 3 of the main text. (**A**) Similar to Fig. 3D, but for trouser and coat concepts. (**B**) Similar to Fig. 3F, but for trouser and coat concepts. (**C**–**E**) Results for cues that combine active neurons from both MF and PP encodings. (**C**) Combined cues formed by the neuron-wise or operation. (**D**) Similar to Fig. 3D, F, but for combined cues. (**E**) Similar to Fig. 3E, G, but for combined cues.

*(Continued on the next page.)*

**Figure S3**: *(Continued from the previous page.)*
(**F**) Overlaps of retrieved patterns over a wide range of network parameters. MF examples and PP concepts are retrieved at high and low threshold, respectively, optimized by grid search. Cue inaccuracy is the fraction of randomly chosen neurons in the target pattern whose activity is flipped to form the cue. Cue incompleteness is the fraction of randomly chosen active neurons in the target pattern which are inactivated to form the cue. Dendritic nonlinearity $\eta$ introduces nonlinear summation between MF patterns $\mathbf{x}^{\mathrm{MF}}_{\mu\nu}$ and PP patterns $\mathbf{x}^{\mathrm{PP}}_{\mu\nu}$ by adding a term $\eta x^{\mathrm{MF}}_{\mu\nu i} x^{\mathrm{PP}}_{\mu\nu i}$ to Eq. 8 of the main text. Negative and positive $\eta$ correspond to sublinear and superlinear regimes, respectively. For PP pattern strength $\gamma = 0.04$, $0.08$, $0.12$, $0.16$, and $0.20$, we respectively use example loads $s = 400$, $120$, $80$, $50$, and $20$; otherwise, $s = 100$. Points represent means over 4 networks with 15 cues tested in each. (**G**, **H**) Similar to Fig. 3H, I, but for different MF pattern densities $a_{\mathrm{MF}}$. PP patterns have correlation $\rho_{\mathrm{PP}} = 0.04$.

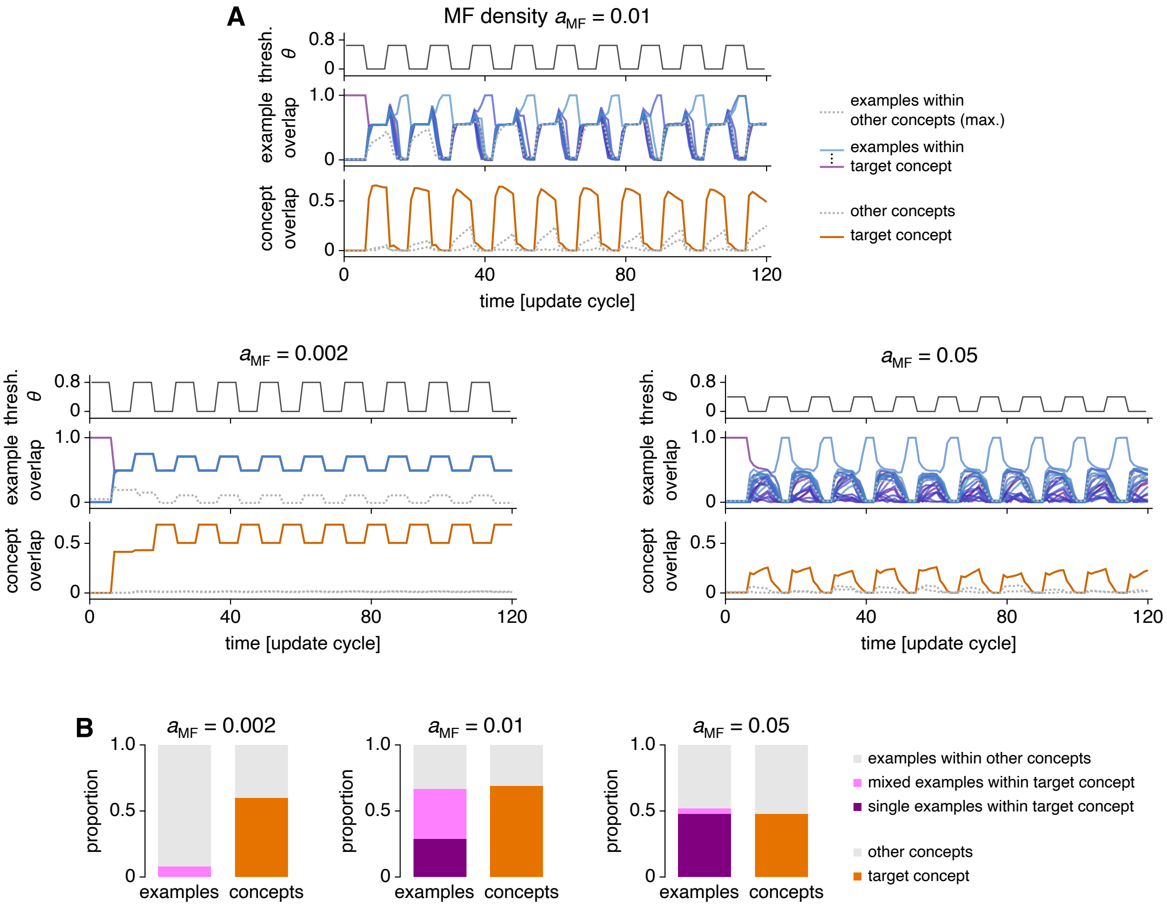


**Figure S4**: Extended results for Fig. 4 of the main text. We use random MF and PP patterns instead of FashionMNIST encodings. (**A**) Similar to Fig. 4A. (**B**) Similar to Fig. 4C. For each scenario in **B**, 10 cues are tested in each of 10 networks. In all networks, 3 concepts are used, MF patterns have correlation 0, and PP patterns have density 0.5 and correlation 0.16. For MF densities $a_{\mathrm{MF}} = 0.002$, $0.01$, and $0.05$, we store 5, 10 and 20 patterns per concept, respectively.

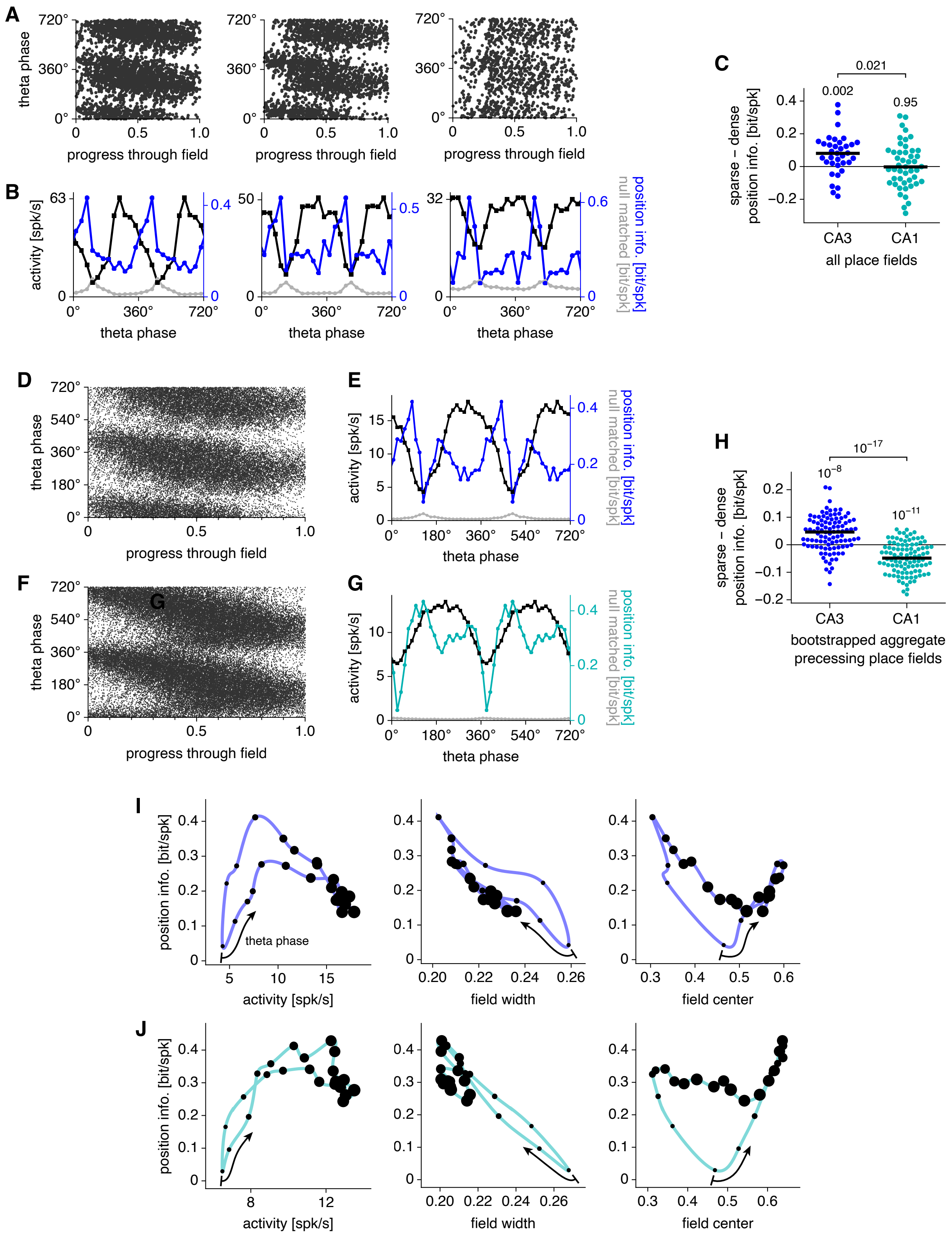


**Figure S5**: Extended results for Fig. 5 of the main text. (**A**–**C**) Additional single-neuron results. (**A**) Similar to Fig. 5F, but for three additional precessing fields from CA3. (**B**) Similar to Fig. 5G, but for the fields in **A**.

*(Continued on the next page.)*

**Figure S5**: *(Continued from the previous page.)*
(**C**) Similar to Fig. 5L, but comparing place fields in CA3 with those in CA1. Numbers indicate $p$-values calculated by two-tailed Wilcoxon signed-rank tests for each population by the two-tailed Mann-Whitney $U$ test for the comparison between them. (**D**–**J**) Results for aggregate fields formed by accumulating spikes across phase precessing place fields. (**D**) Spikes aggregated across 57 CA3 place fields. (**E**) Activity (black), raw position information per spike (blue), and mean null-matched position information (gray) by theta phase for the aggregate field in **D**. (**F**, **G**) Similar to **D**, **E**, but for 55 CA1 place fields. (**H**) Similar to **C**, but comparing 100 bootstrap subsamples per region of 1000 spikes from the aggregate fields in **D** and **F**. (**I**) Parametric plots of features of the aggregate CA3 field in **D** with respect to theta phase. Field width and field center are respectively the standard deviation and mean of progress values over spikes. Each point represents one theta phase and its size is proportional to total activity. Arrows start at the phase with lowest activity and point towards increasing phase. (**J**) Similar to **I**, but for the aggregate CA1 field in **F**.

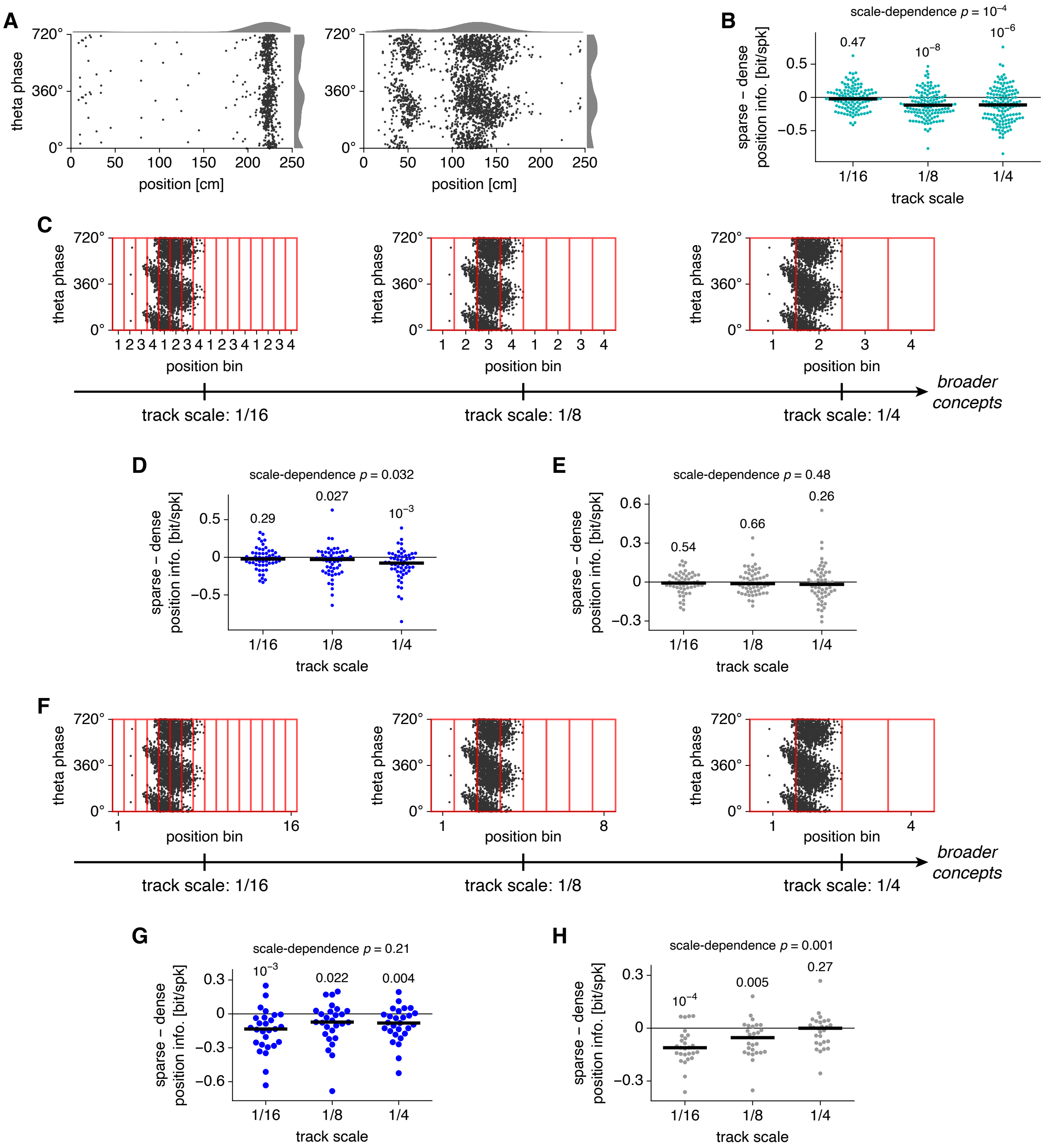


**Figure S6**: Extended results for Fig. 6 of the main text. (**A**) Two additional CA3 place cells along a linear track. For each neuron, spikes are represented by two points at equivalent phases and are accumulated over position (top) and phase (right). (**B**) Similar to Fig. 6E, but for CA1 place cells. (**C**–**E**) Similar to Fig. 6C, E, F, but for an alternative method for binning positions across track scales. (**C**) Four bins are still used for all scales, but bins cycle across the whole track. (**D**) For coarser scales, dense phases convey more position information per spike, as in Fig. 6E. (**E**) Shuffled data exhibit no relationship between position information and theta phase across track scales, as in Fig. 6F. (**F**–**H**) Similar to Fig. 6C, E, F, but for a third method for binning positions across track scales. (**F**) Different numbers of bins are used across scales. (**G**) At all scales, dense phases convey more position information per spike, differing from Fig. 6E. (**H**) Shuffled data exhibit a relationship between position information and theta phase for finer scales, even after sparsity correction, which differs from Fig. 6F and invalidates this binning method.

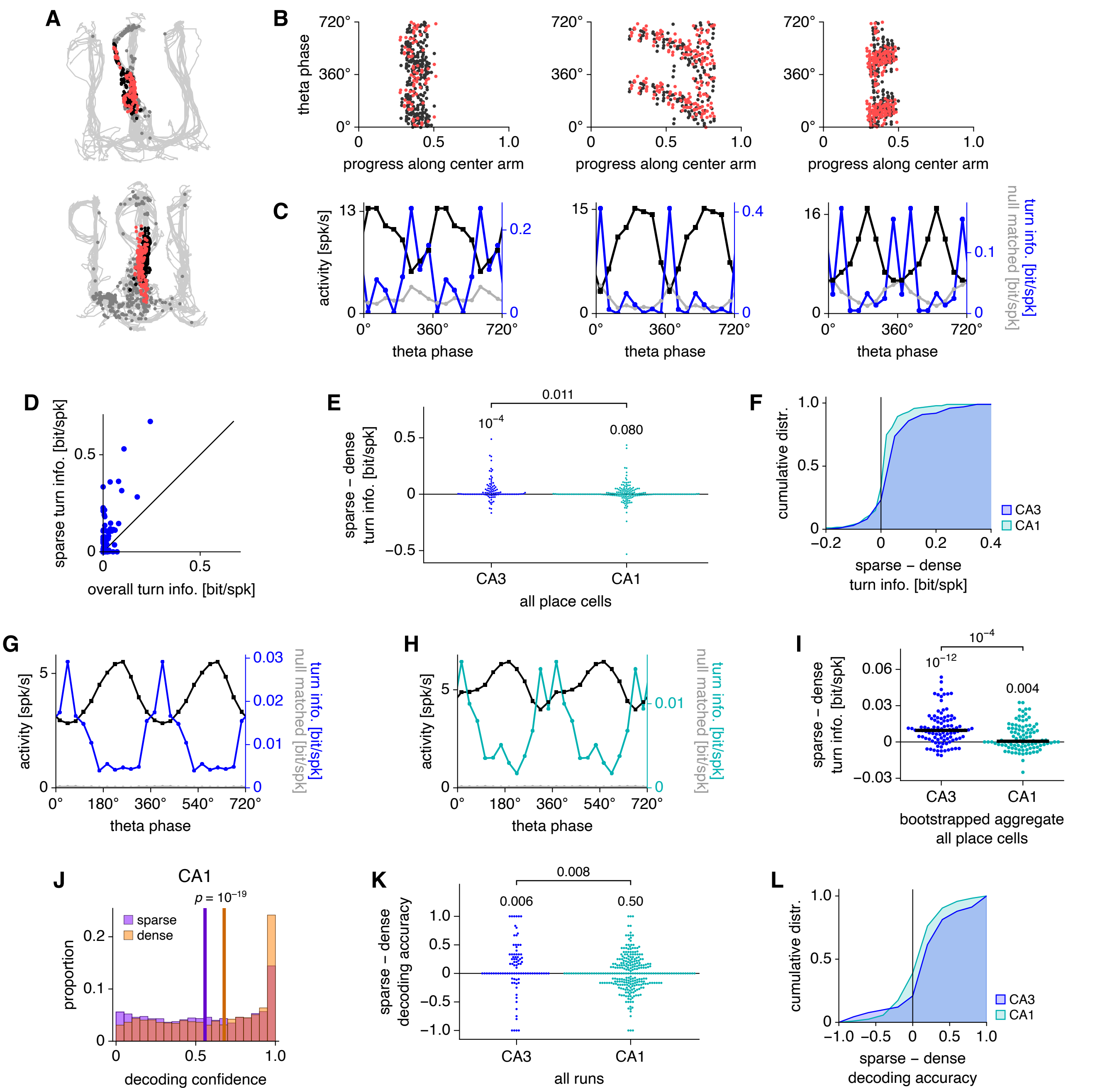


**Figure S7**: Extended results for Fig. 7 of the main text. (**A**–**I**) Additional single-neuron results. (**A**) Spikes from Fig. 7C (top) and Fig. 7E (bottom) superimposed on the animal trajectory (light gray line) and other spikes (dark gray points). (**B**) Similar to Fig. 7C, E, but for three additional CA3 place cells. (**C**) Similar to Fig. 7D, F, but for the fields in **B**. (**D**) Average turn information per spike conveyed by CA3 place cells over sparse theta phases and over all phases. Each point represents one neuron. Note that many neurons convey close to zero overall turn information but convey substantial sparse turn information. (**E**) Similar to Fig. 7G, but comparing place cells in CA3 with those in CA1. Numbers indicate $p$-values calculated by two-tailed Wilcoxon signed-rank tests for each population by the two-tailed Mann-Whitney $U$ test for the comparison between them. (**F**) Cumulative distribution functions for values in **E**. (**G**) Activity (black), raw position information per spike (blue), and mean null-matched position information (gray) by theta phase for spikes aggregated across 98 CA3 place cells. For each place cell, the turn direction with higher activity across all phases is identified. Aggregation is performed by collecting spikes corresponding to more active turn directions and those corresponding to less active directions. (**H**) Similar to **G**, but for 187 CA1 place cells. (**I**) Similar to **E**, but comparing 100 bootstrap subsamples per region of 1000 spikes from the aggregates analyzed in **G** and **H**.

*(Continued on the next page.)*

**Figure S7**: *(Continued from the previous page.)*
(**J**–**L**) Additional Bayesian population decoding results. (**J**) Similar to Fig. 7J, but for CA1 place cells. (**K**) Similar to Fig. 7K, but comparing runs encoded by CA3 with those encoded by CA1. (**L**) Cumulative distribution functions for values in **K**.

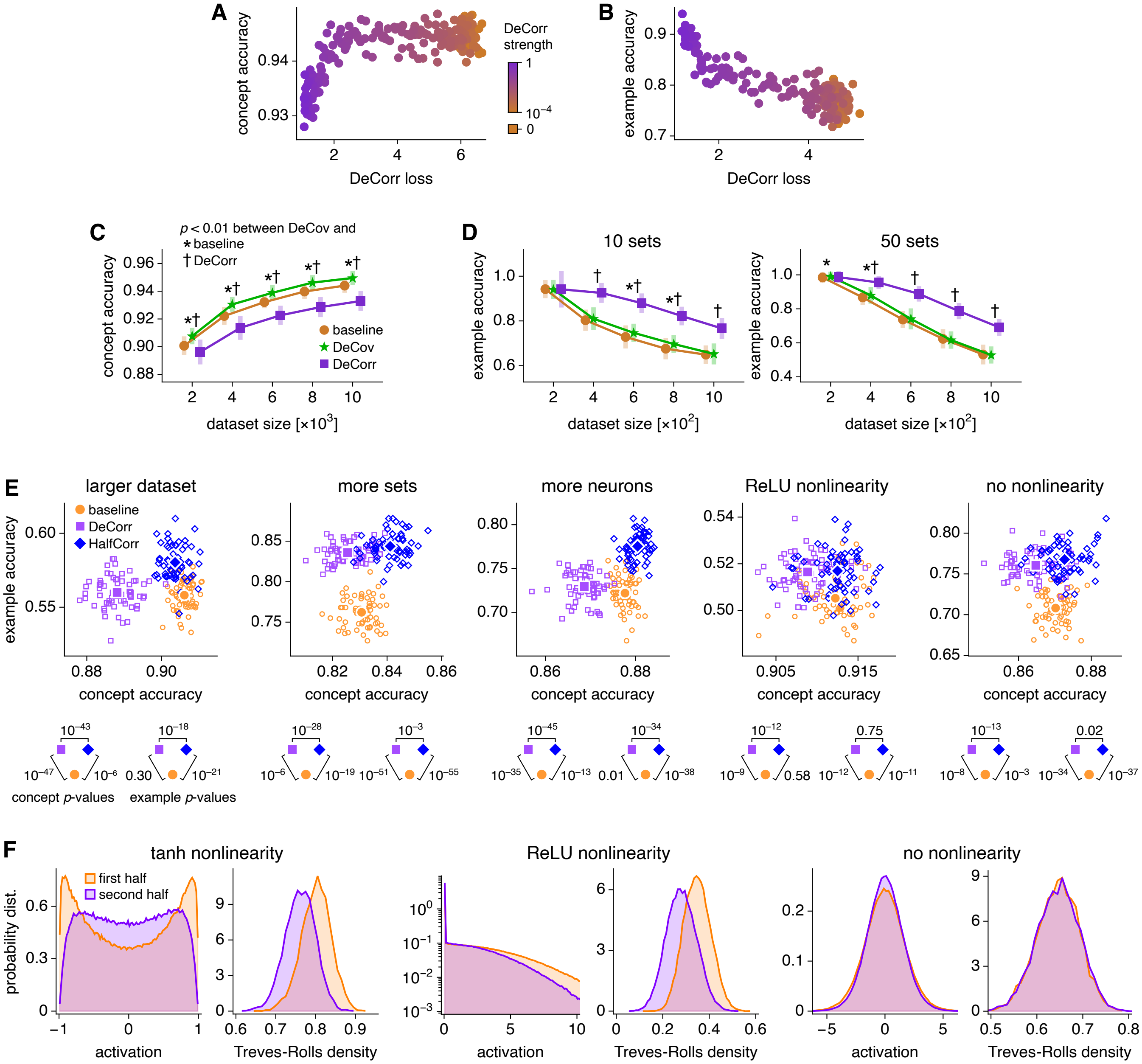


**Figure S8**: Extended results for Fig. 8 of the main text. (**A**–**D**) Additional results for the single-task architecture in Fig. 8B. (**A**, **B**) Concept and example accuracies as functions of DeCorr loss for 192 networks trained with various strengths of the DeCorr loss function. Increasing DeCorr strength decreases the final DeCorr loss, decreases concept accuracy, and increases example accuracy. Dataset sizes are respectively $10\,000$ and $500$ in **A** and **B**, and 100 sets are used in **B**. (**C**, **D**) Similar to Fig. 8E, F, but including networks trained with the DeCov loss function developed by Michael Cogswell and colleagues (Cogswell et al., 2015). Unlike DeCorr, DeCov improves concept accuracy and does not substantially improve example accuracy compared to baseline. (**E**, **F**) Additional results for the multitask architecture in Fig. 8G. (**E**) Similar to Fig. 8I, but for different conditions. In each condition, HalfCorr networks exhibit the best combined performance. From left to right: dataset size of 3000 instead of 1000; 50 sets instead of 10; 500 neurons in each hidden layer instead of 100; ReLU activation function in each hidden layer instead of tanh; and linear activation in the second hidden layer and ReLU activation function in the first hidden layer, which makes the network equivalent to a single-layer perception. (**F**) Activation properties within the final hidden layer of HalfCorr networks with various activation functions described in **E**. Except for the linear activation case, the second, decorrelated half of the layer is sparser than the first, correlated half. Values elicited by 1000 train images in each of 8 trained networks. Treves-Rolls density is the *sparsity* defined in Treves and Rolls (1991) and is computed with the absolute value of activations as in Willmore and Tolhurst (2001).
