## Supplementary Methods for "Distinguishing examples while building concepts in hippocampal and artificial networks"

#### Contents

**Decorrelation in binary feedforward networks** **1**

**CA3 model with random binary patterns** **8**

**CA3 model behavior during oscillating threshold** **8**

**Experimental data preprocessing** **9**

**Experimental data aggregate analysis** **10**

**References** **10**

#### Decorrelation in binary feedforward networks

##### Network architecture

We explore how the correlation of binary activity patterns changes when activity is propagated from one network to another. The two networks are termed presynaptic and postsynaptic. They have sizes $N_{\text{pre}}$ and $N_{\text{post}}$. The presynaptic network exhibits activity patterns $x_{\nu i}^{\text{pre}} \in \{0, 1\}$, where $\nu = 1, \ldots, s$ indexes patterns and $i = 1, \ldots, N_{\text{pre}}$ indexes neurons.

$\mathsf{W}$ is the connectivity matrix from the presynaptic network to the postsynaptic network. For simplicity, we consider binary synaptic weights, so $W_{ij} \in \{0, 1\}$. The postsynaptic patterns $x_{\nu i}^{\text{post}} \in \{0, 1\}$ are determined by a simple threshold operation:

$$x_{\nu i}^{\text{post}} = \Theta\Big[\sum_j W_{ij} x_{\nu j}^{\text{pre}} - \theta\Big], \tag{S1}$$

where $\Theta$ is the Heaviside step function and $\theta$ is the activity threshold. We can rewrite Eq. S1 as

$$x_{\nu i}^{\text{post}} = \Theta\big[g_{\nu i} - \theta\big], \quad \text{where} \quad g_{\nu i} = \sum_j W_{ij} x_{\nu j}^{\text{pre}}. \tag{S2}$$

*

Here, $g_{\nu i}$ is the synaptic input onto postsynaptic neuron $i$ for pattern $\nu$.

We characterize these patterns, either presynaptic or postsynaptic, by their density

$$a = \langle x_{\nu i} \rangle \tag{S3}$$

and the correlation per neuron between two different patterns $\nu \neq \omega$

$$\rho = \frac{\langle x_{\nu i} x_{\omega i} \rangle - \langle x_{\nu i} \rangle \langle x_{\omega i} \rangle}{\langle x_{\nu i} x_{\nu i} \rangle - \langle x_{\nu i} \rangle \langle x_{\nu i} \rangle} = \frac{\langle x_{\nu i} x_{\omega i} \rangle - a^2}{a(1-a)}. \tag{S4}$$

The angle brackets indicate averages over patterns and neurons.

We now assume that the activity patterns and the connectivity matrix are generated via random processes. To be explicit, we will write $X_{\nu i}^{\mathrm{pre}}$ as the random variable for the activity of presynaptic neuron $i$ in pattern $\nu$; the same capitalization applies to the postsynaptic activities $X_{\nu i}^{\mathrm{post}}$ and inputs $G_{\nu i}$. Lowercase letters represent samples of these variables.

We assume that each $X_{\nu i}^{\mathrm{pre}}$ is an identically distributed random variable. We assume that the $W_{ij}$'s are independent and identically distributed (iid) Bernoulli random variables with parameter $u$:

$$W_{ij} \sim \mathrm{Ber}(u). \tag{S5}$$

In this case, each $X_{\nu i}^{\mathrm{post}}$ is also an identically distributed random variable.

We can then write expressions for density and correlation as

$$\begin{aligned} a &= \mathrm{E}[X_{\nu i}] \\ \rho &= \frac{\mathrm{E}[X_{\nu i} X_{\omega i}] - a^2}{a(1-a)}. \end{aligned} \tag{S6}$$

These are population values; Eqs. S3 and S4 indicate the sample estimates and should be written as $\hat{a}$ and $\hat{\rho}$, but we will ignore this distinction.

#### Generating presynaptic activity patterns

For mathematical tractability, we will enforce density on a per-pattern basis; that is, each presynaptic pattern has the same number of active neurons:

$$n \equiv N_{\mathrm{pre}} a_{\mathrm{pre}} = \sum_i X_{\nu i}^{\mathrm{pre}}. \tag{S7}$$

We generate correlated patterns obeying this restriction as follows:

1. We create a concept pattern $\bar{\mathbf{x}}^{\mathrm{pre}} \in \{0,1\}^{N_{\mathrm{pre}}}$ by randomly choosing $n$ neurons to be 1 and the rest to be 0. Let $\mathcal{S}$ be the set of all active neurons (value 1) and its complement $\mathcal{S}^{\mathrm{c}}$ be the set of all inactive neurons (value 0).

2. To create each example pattern $\mathbf{x}_{\nu}^{\mathrm{pre}}$, we randomly select a fraction $d$ of neurons in $\mathcal{S}$ and set them to 0. We then randomly select the same number $nd$ of neurons in $\mathcal{S}^{\mathrm{c}}$ and set them to 1.

Now we calculate the correlation $\rho_{\mathrm{pre}}$ between the patterns $\mathbf{X}_{\nu}^{\mathrm{pre}}$ generated by this method. To do so, we investigate the distribution of $M$, a random variable for the number of active neurons common to two different patterns. We note that

$$M = M_{\mathcal{S}} + M_{\mathcal{S}^{\mathrm{c}}}, \tag{S8}$$

where $M_{\mathcal{S}}$ is the number of neurons in $\mathcal{S}$ that remain active in both patterns, and $M_{\mathcal{S}^{\mathrm{c}}}$ is the number of neurons in $\mathcal{S}^{\mathrm{c}}$ that are activated in both patterns. These numbers have hypergeometric distributions:

$$\begin{aligned} M_{\mathcal{S}} &\sim \mathrm{Hyp}\big[n, n(1-d), n(1-d)\big], \\ M_{\mathcal{S}^{\mathrm{c}}} &\sim \mathrm{Hyp}\big[N_{\mathrm{pre}} - n, nd, nd\big]. \end{aligned} \tag{S9}$$

We can then calculate

$$\mathrm{E}[X^{\mathrm{pre}}_{\nu i} X^{\mathrm{pre}}_{\omega i}] = \frac{\mathrm{E}[M]}{N_{\mathrm{pre}}} = \frac{1}{N_{\mathrm{pre}}}\left(\frac{n^2(1-d)^2}{n} + \frac{n^2 d^2}{N_{\mathrm{pre}} - n}\right) = a_{\mathrm{pre}}(1-d)^2 + \frac{a_{\mathrm{pre}}^2 d^2}{1 - a_{\mathrm{pre}}}. \tag{S10}$$

Substituting this expression into Eq. S6, we obtain

$$\begin{aligned} \rho_{\mathrm{pre}} &= \left(\frac{1 - a_{\mathrm{pre}} - d}{1 - a_{\mathrm{pre}}}\right)^2, \\ d &= \big(1 - a_{\mathrm{pre}}\big)\big(1 - \sqrt{\rho_{\mathrm{pre}}}\big). \end{aligned} \tag{S11}$$

Thus, we know the number of activity flips required to produce presynaptic patterns with a given correlation.

#### Identifying the probability distribution of $G_{\nu i}$

Postsynaptic activity patterns are produced by Eq. S2. Written in terms of random variables, it becomes

$$X^{\mathrm{post}}_{\nu i} = \Theta\big[G_{\nu i} - \theta\big], \quad \text{where} \quad G_{\nu i} = \sum_j W_{ij} X^{\mathrm{pre}}_{\nu j}. \tag{S12}$$

Thus, the statistics of $\mathbf{X}^{\mathrm{post}}_{\nu}$ are determined by $\mathbf{G}_{\nu}$. To calculate second-order statistics such as $\rho_{\mathrm{post}}$, we need to determine the joint distribution of $G_{\nu i}$ and $G_{\omega i}$ for two patterns $\nu \neq \omega$.

To do so, we define $\mathcal{R}_0$ as the set of active neurons that are common to both patterns $\mathbf{X}^{\mathrm{pre}}_{\nu}$ and $\mathbf{X}^{\mathrm{pre}}_{\omega}$. We define $\mathcal{R}_{\nu}$ and $\mathcal{R}_{\omega}$ as sets of neurons only active in patterns $\nu$ and $\omega$, respectively. We can then write

$$\begin{aligned} G_{\nu i} &= \sum_{j \in \mathcal{R}_0} W_{ij} + \sum_{j \in \mathcal{R}_{\nu}} W_{ij} \equiv G_{0i} + \tilde{G}_{\nu i} \\ G_{\omega i} &= \sum_{j \in \mathcal{R}_0} W_{ij} + \sum_{j \in \mathcal{R}_{\omega}} W_{ij} \equiv G_{0i} + \tilde{G}_{\omega i}. \end{aligned} \tag{S13}$$

These sets have cardinalities

$$|\mathcal{R}_0| = M, \quad |\mathcal{R}_{\nu}| = n - M, \quad |\mathcal{R}_{\omega}| = n - M, \tag{S14}$$

where $n$ is given by Eq. S7 and $M$ is given by Eqs. S8 and S9. Since the elements of $\mathsf{W}$ are iid Bernoulli random variables (Eq. S5),

$$\begin{aligned}
\big(G_{0i} \mid M = m\big) &\sim \mathrm{Bin}(m, u),\\
\big(\tilde{G}_{\nu i} \mid M = m\big) &\sim \mathrm{Bin}(n-m, u),\\
\big(\tilde{G}_{\omega i} \mid M = m\big) &\sim \mathrm{Bin}(n-m, u).
\end{aligned} \tag{S15}$$

Since $\mathcal{R}_0$, $\mathcal{R}_\nu$, and $\mathcal{R}_\omega$ are mutually disjoint, $G_{0i}$, $\tilde{G}_{\nu i}$, and $\tilde{G}_{\omega i}$ are mutually independent when conditioned on $M$.

#### Calculating the joint probability density function of $G_{\nu i}$ and $G_{\omega i}$

We take the $N_{\mathrm{pre}} \to \infty$ limit in which the discrete probability distributions for $X_{\nu i}$ and $G_{\nu i}$ are replaced by their continuous limits. First, we present a few points on notation.

1. A continuous random variable $A$ has probability density $f_A(a)$ at value $A = a$. We may simplify it as $f(a)$.
2. Variables $A$ and $B$ have joint probability density $f_{A,B}(a, b)$ at values $A = a$ and $B = b$. We may simplify it as $f(a, b)$.
3. Similarly, the conditional probability density given that the random variable $B$ takes value $b$ is $f_{A|B=b}(a)$. We may simplify it as $f(a|b)$.

We can now write the joint probability density function (pdf) for $G_{\nu i}$ and $G_{\omega i}$. For notational convenience, we will drop the subscript $i$ for all relevant variables. The pdf is

$$\begin{aligned}
f(g_\nu, g_\omega) &= \int \mathrm{d}g_0\, f(g_\nu, g_\omega, g_0)\\
&= \int \mathrm{d}g_0\, f_{\tilde{G}_\nu, \tilde{G}_\omega, G_0}(g_\nu - g_0, g_\omega - g_0, g_0)\\
&= \int \mathrm{d}m \int \mathrm{d}g_0\, f_{\tilde{G}_\nu, \tilde{G}_\omega, G_0}(g_\nu - g_0, g_\omega - g_0, g_0 \mid m) f(m)\\
&= \int \mathrm{d}m \int \mathrm{d}g_0\, f_{\tilde{G}_\nu}(g_\nu - g_0|m) f_{\tilde{G}_\omega}(g_\omega - g_0|m) f(g_0|m) f(m).
\end{aligned} \tag{S16}$$

The second line is obtained using the change-of-variables formula for a joint pdf.

We have expressions for each pdf in the integrand. In the large $N_{\mathrm{pre}}$ limit, the binomial distributions in Eq. S15 can be approximated by normal distributions (dropping the subscript $i$ for notational convenience):

$$\begin{aligned}
\big(G_0 \mid M = m\big) &\sim \mathcal{N}\big[mu, mu(1-u)\big],\\
\big(\tilde{G}_\nu \mid M = m\big) &\sim \mathcal{N}\big[(n-m)u, (n-m)u(1-u)\big],\\
\big(\tilde{G}_\omega \mid M = m\big) &\sim \mathcal{N}\big[(n-m)u, (n-m)u(1-u)\big].
\end{aligned} \tag{S17}$$

Thus, we find

$$
\begin{aligned}
&\int \mathrm{d}g_0\, f_{\tilde{G}_\nu}(g_\nu - g_0|m) f_{\tilde{G}_\omega}(g_\omega - g_0|m) f(g_0|m) \\
&\quad \propto \int \mathrm{d}g_0\, \exp\left[-\frac{\big(g_\nu - g_0 - (n-m)u\big)^2}{2(n-m)u(1-u)}\right] \exp\left[-\frac{\big(g_\omega - g_0 - (n-m)u\big)^2}{2(n-m)u(1-u)}\right] \exp\left[-\frac{(g_0 - mu)^2}{2mu(1-u)}\right] \\
&\quad = \int \mathrm{d}g_0\, \exp\left[-\frac{\left[g_0 - \left(\frac{g_\nu + g_\omega}{2} - (n-m)u\right)\right]^2}{(n-m)u(1-u)}\right] \exp\left[-\frac{\left(\frac{g_\nu - g_\omega}{2}\right)^2}{(n-m)u(1-u)}\right] \exp\left[-\frac{(g_0 - mu)^2}{2mu(1-u)}\right] \\
&\quad \propto \exp\left[-\frac{\left(\frac{g_\nu + g_\omega}{2} - nu\right)^2}{(n+m)u(1-u)}\right] \exp\left[-\frac{\left(\frac{g_\nu - g_\omega}{2}\right)^2}{(n-m)u(1-u)}\right].
\end{aligned}
\tag{S18}
$$

We can write the terms inside the exponential as

$$
\begin{aligned}
&\frac{\left(\frac{g_\nu + g_\omega}{2} - nu\right)^2}{(n+m)u(1-u)} + \frac{\left(\frac{g_\nu - g_\omega}{2}\right)^2}{(n-m)u(1-u)} \\
&\quad = \frac{\left[\frac{(g_\nu - nu) + (g_\omega - nu)}{2}\right]^2}{(n+m)u(1-u)} + \frac{\left[\frac{(g_\nu - nu) - (g_\omega - nu)}{2}\right]^2}{(n-m)u(1-u)} \\
&\quad = \frac{1}{2}\left[\begin{pmatrix} g_\nu - nu & g_\omega - nu \end{pmatrix} \frac{1}{2u(1-u)} \begin{pmatrix} \frac{1}{n+m} + \frac{1}{n-m} & \frac{1}{n+m} - \frac{1}{n-m} \\ \frac{1}{n+m} - \frac{1}{n-m} & \frac{1}{n+m} + \frac{1}{n-m} \end{pmatrix} \begin{pmatrix} g_\nu - nu \\ g_\omega - nu \end{pmatrix}\right] \\
&\quad = \frac{1}{2}(\mathbf{g} - \boldsymbol{\mu}_G)^\top \boldsymbol{\Sigma}_G^{-1} (\mathbf{g} - \boldsymbol{\mu}_G).
\end{aligned}
\tag{S19}
$$

The last expression is written in terms of the variable vector, mean vector, and covariance matrix for $G_\nu$ and $G_\mu$:

$$
\begin{aligned}
\mathbf{g} &= \begin{pmatrix} g_\nu \\ g_\omega \end{pmatrix} \\
\boldsymbol{\mu}_G &= \begin{pmatrix} nu \\ nu \end{pmatrix} = \begin{pmatrix} N_{\mathrm{pre}} a_{\mathrm{pre}} u \\ N_{\mathrm{pre}} a_{\mathrm{pre}} u \end{pmatrix} \\
\boldsymbol{\Sigma}_G &= \begin{pmatrix} nu(1-u) & mu(1-u) \\ mu(1-u) & nu(1-u) \end{pmatrix} = \sigma_G^2 \begin{pmatrix} 1 & \rho_G \\ \rho_G & 1 \end{pmatrix},
\end{aligned}
\tag{S20}
$$

where the covariance and correlation are

$$
\sigma_G^2 = N_{\mathrm{pre}} a_{\mathrm{pre}} u(1-u) \quad \text{and} \quad \rho_G = \frac{m}{N_{\mathrm{pre}} a_{\mathrm{pre}}}.
\tag{S21}
$$

Combining Eqs. S16, S18, and S19, we obtain

$$
f(g_\nu, g_\omega) \propto \int \mathrm{d}m\, \exp\left[-\frac{1}{2}(\mathbf{g} - \boldsymbol{\mu}_G)^\top \boldsymbol{\Sigma}_G^{-1} (\mathbf{g} - \boldsymbol{\mu}_G)\right] f(m).
\tag{S22}
$$

Now we consider $M = M_{\mathcal{S}} + M_{\mathcal{S}^c}$ (Eq. S8). From Eq. S9, we see that $M_{\mathcal{S}}$ and $M_{\mathcal{S}^c}$ have means and variances

$$\begin{aligned}
\mu_{\mathcal{S}} &= n(1-d)^2 && = N_{\text{pre}} a_{\text{pre}} (1-d)^2 \\
\sigma_{\mathcal{S}}^2 &= n d^2 (1-d)^2 && = N_{\text{pre}} a_{\text{pre}} d^2 (1-d)^2 \\
\mu_{\mathcal{S}^c} &= \frac{n^2 d^2}{N_{\text{pre}} - n} && = N_{\text{pre}} \frac{a_{\text{pre}}^2 d^2}{1 - a_{\text{pre}}} \\
\sigma_{\mathcal{S}^c}^2 &= \frac{n^2 d^2 \big(N_{\text{pre}} - n(1+d)\big)^2}{(N_{\text{pre}} - n)^3} && = N_{\text{pre}} \frac{a_{\text{pre}}^2 d^2 (1 - a_{\text{pre}} - a_{\text{pre}} d)^2}{(1 - a_{\text{pre}})^3}.
\end{aligned} \tag{S23}$$

Note that the flip fraction $d$ can be expressed in terms of $a_{\text{pre}}$ and $\rho_{\text{pre}}$ via Eq. S11.

As $N_{\text{pre}} \to \infty$, the hypergeometric random variables $M_{\mathcal{S}}$ and $M_{\mathcal{S}^c}$ approach normal distributions with the means and variances in Eq. S23. Thus, their distributions become sharply peaked around their means, and we can approximate the pdf of $M$ by a delta-function at its mean:

$$\begin{aligned}
f(m) &\to \delta(m - \mu_M), \quad \text{where} \\
\mu_M &= N_{\text{pre}} a_{\text{pre}} (1-d)^2 + N_{\text{pre}} \frac{a_{\text{pre}}^2 d^2}{1 - a_{\text{pre}}} \\
&= N_{\text{pre}} a_{\text{pre}} (a_{\text{pre}} + \rho_{\text{pre}} - a_{\text{pre}} \rho_{\text{pre}}).
\end{aligned} \tag{S24}$$

We now have our final expression for the joint pdf of $G_{\nu i}$ and $G_{\omega}$. Reintroducing the neural index $i$ and the normalization factor, Eq. S22 becomes

$$f(g_{\nu i}, g_{\omega i}) = \frac{1}{2\pi \sqrt{\det \mathbf{\Sigma}_G}} \exp\left[ -\frac{1}{2} (\mathbf{g}_i - \boldsymbol{\mu}_G)^\top \mathbf{\Sigma}_G^{-1} (\mathbf{g}_i - \boldsymbol{\mu}_G) \right], \tag{S25}$$

where

$$\mathbf{g}_i = \begin{pmatrix} g_{\nu i} \\ g_{\omega i} \end{pmatrix}, \qquad \boldsymbol{\mu}_G = \begin{pmatrix} \mu_G \\ \mu_G \end{pmatrix}, \qquad \mathbf{\Sigma}_G = \sigma_G^2 \begin{pmatrix} 1 & \rho_G \\ \rho_G & 1 \end{pmatrix} \tag{S26}$$

and

$$\mu_G = N_{\text{pre}} a_{\text{pre}} u, \qquad \sigma_G^2 = N_{\text{pre}} a_{\text{pre}} u(1-u), \qquad \rho_G = a_{\text{pre}} + \rho_{\text{pre}} - a_{\text{pre}} \rho_{\text{pre}}. \tag{S27}$$

Figure S1 shows a plot of the joint pdf $f(g_{\nu i}, g_{\omega i})$ along with a histogram obtained through numerical simulation. The theoretical formula Eq. S25 agrees very well with the numerical data.

#### Integrating the joint probability density function to obtain $a_{\text{post}}$ and $\rho_{\text{post}}$

With the joint pdf for $G_{\nu i}$ and $G_{\omega i}$ (Eq. S25), we can compute the postsynaptic pattern density $a_{\text{post}}$ and correlation $\rho_{\text{post}}$ using Eqs. S6 and S12. According to Eq. S12, $X_{\nu i}^{\text{post}}$ acts as an indicator random variable for $G_{\nu i} > \theta$, and the product $X_{\nu i}^{\text{post}} X_{\omega i}^{\text{post}}$ acts as an indicator random variable for $G_{\nu i} > \theta \cap G_{\omega i} > \theta$. Thus,

$$\mathrm{E}[X_{\nu i}^{\text{post}}] = P(G_{\nu i} > \theta) \quad \text{and} \quad \mathrm{E}[X_{\nu i}^{\text{post}} X_{\omega i}^{\text{post}}] = P(G_{\nu i} > \theta \cap G_{\omega i} > \theta). \tag{S28}$$

We can calculate

$$\begin{aligned}
\mathrm{E}[X_{\nu i}^{\text{post}}] &= \int_{\theta}^{\infty} \mathrm{d}g_{\nu i} \int_{-\infty}^{\infty} \mathrm{d}g_{\omega i}\, f(g_{\nu i}, g_{\omega i}) \\
&= \int_{\theta}^{\infty} \mathrm{d}g_{\nu i}\, \frac{1}{\sqrt{2\pi}\sigma_G} \exp\!\left[-\frac{(g_{\nu i}-\mu_G)^2}{2\sigma_G^2}\right] \\
&= \frac{1}{2} \operatorname{erfc} \frac{\theta - \mu_G}{\sqrt{2}\sigma_G}.
\end{aligned} \tag{S29}$$

Thus, the postsynaptic pattern density is immediately

$$a_{\text{post}} = \mathrm{E}[X_{\nu i}^{\text{post}}] = \frac{1}{2} \operatorname{erfc} \frac{\phi}{\sqrt{2}}, \quad \text{where} \quad \phi = \frac{\theta - \mu_G}{\sigma_G}. \tag{S30}$$

The rescaled threshold $\phi$ is the standardized version of $\theta$.

We next need to calculate

$$\begin{aligned}
\mathrm{E}[X_{\nu i}^{\text{post}} X_{\omega i}^{\text{post}}] &= \int_{\theta}^{\infty} \mathrm{d}g_{\nu i} \int_{\theta}^{\infty} \mathrm{d}g_{\omega i}\, f(g_{\nu i}, g_{\omega i}) \\
&= \frac{1}{2\pi\sqrt{\det \boldsymbol{\Sigma}_G}} \int_{\theta}^{\infty} \mathrm{d}g_{\nu i} \int_{\theta}^{\infty} \mathrm{d}g_{\omega i}\, \exp\!\left[-\frac{1}{2}(\mathbf{g}_i - \boldsymbol{\mu}_G)^{\top} \boldsymbol{\Sigma}_G^{-1} (\mathbf{g}_i - \boldsymbol{\mu}_G)\right].
\end{aligned} \tag{S31}$$

By standardizing the variables of integration with $h_{\nu i} = (g_{\nu i} - \mu_G)/\sigma_G$, this integral can be expressed in terms of the standard bivariate normal:

$$\mathrm{E}[X_{\nu i}^{\text{post}} X_{\omega i}^{\text{post}}] = \frac{1}{2\pi\sqrt{1-\rho_G^2}} \int_{\phi}^{\infty} \mathrm{d}h_{\nu i} \int_{\phi}^{\infty} \mathrm{d}h_{\omega i}\, \exp\!\left[-\frac{h_{\nu i}^2 + h_{\omega i}^2 - 2\rho_G h_{\nu i} h_{\omega i}}{2(1-\rho_G^2)}\right]. \tag{S32}$$

This double integral cannot be evaluated in closed form, but we can reduce it to a single integral (Owen, 1956):

$$\mathrm{E}[X_{\nu i}^{\text{post}} X_{\omega i}^{\text{post}}] = \Gamma[\phi, \rho_G] \equiv \frac{1}{2\pi} \int_{\arccos \rho_G}^{\pi} \mathrm{d}\psi\, \exp\!\left[-\frac{\phi^2}{1 + \cos\psi}\right]. \tag{S33}$$

Therefore, the expression for the postsynaptic correlation follows:

$$\rho_{\text{post}} = \frac{\Gamma[\phi, \rho_G] - a_{\text{post}}^2}{a_{\text{post}}(1 - a_{\text{post}})}, \tag{S34}$$

where $a_{\text{post}}$ can be expressed in terms of the standardized threshold $\phi$ with Eq. S30. On the other hand, we can stipulate a desired $a_{\text{post}}$ and then recover $\phi$ and $\rho_{\text{post}}$ with

$$\begin{aligned}
\phi &= \sqrt{2} \operatorname{erfc}^{-1}(2a_{\text{post}}), \\
\rho_{\text{post}} &= \frac{\Gamma\!\left[\sqrt{2} \operatorname{erfc}^{-1}(2a_{\text{post}}),\, a_{\text{pre}} + \rho_{\text{pre}} - a_{\text{pre}}\rho_{\text{pre}}\right] - a_{\text{post}}^2}{a_{\text{post}}(1 - a_{\text{post}})}.
\end{aligned} \tag{S35}$$

This is Eq. 1 of the main text. Figure 2E shows that this formula for the postsynaptic correlation agrees well with values obtained through numerical simulation across a variety of parameter values.

#### Exploring $\rho_{\text{post}}$ as a function of $a_{\text{pre}}$, $a_{\text{post}}$, and $\rho_{\text{pre}}$

In Fig. S2B, we plot $\rho_{\text{post}}$ as a function $a_{\text{pre}}$ and $a_{\text{post}}$ for various $\rho_{\text{pre}}$. We see that decorrelation ($\rho_{\text{post}} < \rho_{\text{pre}}$) occurs when $a_{\text{post}}$ is low. Thus, a downstream (postsynaptic) network with many neurons but low activity naturally decorrelates patterns of the upstream (presynaptic) network. The low activity can be achieved by low connectivity $u$ or a high threshold $\theta$; Eq. S35 does not differentiate between the two.

Figure S2C demonstrates that if presynaptic patterns are sparse and decorrelated (lower left corner of the plot), postsynaptic patterns also exhibit low correlation even if they are denser. Thus, once patterns are sparsified and decorrelated, they will remain decorrelated for subsequent feedforward layers. Note that this panel also shows the symmetry in interchanging $a_{\text{pre}} \leftrightarrow \rho_{\text{pre}}$ present in Eq. S35.

### CA3 model with random binary patterns

We use a network size of $N_{\text{CA3}} = 10\,000$. We generate MF example patterns $\mathbf{x}_{\mu\nu}^{\text{MF}}$ with desired density $a_{\text{MF}}$ and correlation 0 by randomly activating $N_{\text{CA3}} a_{\text{MF}}$ neurons. We generate PP concept patterns $\bar{\mathbf{x}}_{\mu}^{\text{PP}}$ with desired density 0.5 by randomly activating each neuron with probability 0.5. We then generate PP example patterns $\mathbf{x}_{\mu\nu}^{\text{PP}}$ with desired correlation $\rho_{\text{PP}}$ by randomly flipping each concept neuron with probability $(1 - \sqrt{\rho_{\text{PP}}})/2$. Simulations are initiated without cue noise and a sharp activation threshold ($\beta \to \infty$) to assess the best possible network performance.

To calculate capacities in Fig. 3H, I, we use a higher strength of PP inputs $\zeta = 0.2$ to make the capacity values more computationally accessible. For a given load of concepts per neuron, we perform a grid search over the load of examples per concept using 8 networks per load and testing 20 cues in each network. Each cue is identical to its target pattern. We search for the load at which the average overlap crosses a threshold, which is 1/2, $(1 + \rho_{\text{PP}})/2$, and $(1 + \sqrt{\rho_{\text{PP}}})/2$ for MF examples, PP examples, and PP concepts, respectively, to account for the positive overlap of off-target PP patterns if they are correlated (Kang and Toyoizumi, 2023). For MF examples, we explore activity thresholds $\theta'$ between 0.43 and 0.85 and use the value that maximizes average overlap. For PP examples and concepts, we set the activity threshold $\theta'$ to 0.

### CA3 model behavior during oscillating threshold

The oscillation analysis in Fig. 4C characterizes network behavior between update cycles 60 and 120. Consider a single oscillation cycle. For example-related behavior, we consider its high-threshold half. Let $m_1(t)$ and $m_2(t)$ be the largest and second-largest overlaps within the target concept at time $t$, and let $m_0(t)$ be the largest overlap within other concepts. If $m_1(t) > 0.8$, $m_1(t) > 2 \cdot m_2(t)$, and $m_1(t) > 2 \cdot m_0(t)$ at any time, the behavior of the oscillation cycle is categorized as *single examples within a target concept*. If $m_1(t) > 0.8$, $m_1(t) < 2 \cdot m_2(t)$, and $m_1(t) > 2 \cdot m_0(t)$, the behavior is categorized as *mixed examples within a target concept*. If at least half of the oscillation cycles receive a certain categorization, it is considered the network behavior. Otherwise, the network behavior is *examples within other concepts*. For concept-related behavior, we consider the low-threshold half of each oscillation cycle. Let $m_1(t)$ be the overlap with the target concept and $m_0(t)$ be the largest overlap with other concepts. If $m_1(t) > 0.1$ and $m_1(t) > 2 \cdot m_0(t)$ at any time, the behavior of the oscillation cycle is categorized as *target concept* (for the random binary patterns in Fig. S4B, we use $m_1(t) > 0.2$ instead). If at least half of the oscillation cycles receive this categorization, it is considered the network behavior. Otherwise, the network behavior is *other concepts*.

### Experimental data preprocessing

In all experimental analyses, we consider each traveling direction separately. Thus, each recorded neuron effectively yields two neurons in our analyses with their own spikes and trajectory occupancies.

Linear track data from the CRCNS hc-3 dataset is used to produce the results in Figs. 5 and 6 (Mizuseki et al., 2013). For CA3, we use all linear track sessions from rats ec013, ec016, gor, and vvp with CA3 neurons recorded, and for CA1, we use all linear track sessions between 761 and 882 from rat ec013. For rats gor and vvp, the size of the linear track was changed during recording, and we only consider data before the change. Animal positions are taken to be the mean of the two LED lights on the microdrive. The track axis is taken to be the first principal component of sampled positions. Animal velocities are differences between tracked position samples divided by the sampling rate and smoothed with a Gaussian filter with standard deviation 0.1 s.

We use the recommended quality criteria involving eDist, RefracRatio, and RefracViol when selecting units. Since we are interested in the theta oscillation, we only consider spikes occurring during locomotion with speed greater than 10 cm/s. We require units to have at least 50 spikes occurring within the central 70% of the track to avoid neurons whose behavior may be dominated by boundary effects. To determine the theta signal for each unit, we identify the tetrode from which it was recorded and average the LFP over all channels on the tetrode. This signal is bandpass-filtered between 6–10 Hz, and the complex argument of its Hilbert transform is the local theta phase.

To extract place fields for Fig. 5, we first discretize track positions with 1 cm bins. For a given place cell, we first compute the activity across all theta phases as a function of position and apply a Gaussian filter whose standard deviation is 0.01 times the track length. We find activity peaks whose maximum is 0.6 standard deviations above the mean; if the activity exhibits multiple peaks while remaining above this threshold, the largest is chosen. We then find the closest flanking positions where the activity falls below 0.2 times the peak value. The region in between is the place field. If two place fields overlap, they are divided at the activity minimum located between their peaks.

W-maze data from the CRCNS hc-6 dataset is used to produce the results in Fig. 7 (Karlsson et al., 2015). We use all run sessions from rats bon, con, dud, fra, mil, and ten. We remove 2 sessions from dud and 3 sessions from fra without position samples in one of the side arms. Animal positions and velocities are taken directly as the 30 Hz-interpolated samples from the dataset. We only consider spikes occurring during locomotion with speed greater than 5 cm/s. We require units to have at least 30 spikes occurring within the center arm. Spike theta phases are taken directly from the dataset.

To extract arm identity from the position samples, we first linearly rescale both position coordinates to span from 0 to 1. We define a maze skeleton consisting of lines that represent the left arm from $(0.1, 0.1)$ to $(0.1, 1)$, the center arm from $(0.5, 0.1)$ to $(0.5, 1)$, the right arm from $(0.9, 0.1)$ to $(0.9, 1)$, and the base from $(0.1, 0.1)$ to $(0.9, 0.1)$. We then fit transformations of this skeleton to the position samples, allowing for stretching along the first coordinate, rotation about its center, and translations. The fit is performed by minimizing the total squared distance between each sampled position and its closest point on the transformed skeleton. Empirically, this process yields excellent fitting without any need for manual intervention. Then, each position sample is assigned to either the left, center, or right arm based on closest distance.

To extract runs along the central arm, we first smooth arm samples by encoding arm identity as a one-hot vector, applying a Gaussian filter with standard deviation 0.2 s, and identifying the largest element in each sample. We then consider each span of central-arm samples. We find the times within the span at which the animal crosses scaled positions 0.4 and 0.7, where 0 corresponds to the smallest position value at the base of the maze and 1 corresponds to the largest position value at the far end of the arms. The run duration must be between 0.1 s and 10 s, and we then pad the run by 0.5 s at both ends. Outward runs cross scaled position 0.7 before 0.4, and the subsequent arm identity is used to determine the future turn direction. Inward runs cross scaled position 0.4 before 0.7, and the previous arm identity is used to determine the past turn direction.

### Experimental data aggregate analysis

The aggregate fields in Fig. S5D–J are formed from phase-precessing place fields. We do not enforce a minimum spike count or ensure theta modulation on a single-neuron basis. In total, we collect 19 678 spikes from 57 CA3 place fields and 29 664 spikes from 55 CA1 place fields. We perform bootstrapping by sampling with replacement 1000 spikes at a time. For each subsample, we bin spikes into 10 progress bins and phase bins of width 15°.

The aggregate fields in Fig. S7G–I are formed from place cells with at least 30 spikes within the central arm. We do not ensure theta modulation on a single-neuron basis. For each neuron, we identify the turn directions with higher and lower activities, and collect spikes occurring during each condition. In total, we collect 47 196 spikes from 331 CA3 place cells and 72 461 spikes from 436 CA1 place cells. We perform bootstrapping by sampling with replacement 1000 spikes at a time. For each subsample, we bin spikes into 2 directions (more active and less active) and phase bins of width 30°.
